# Cognitive Resilience in Aging Degus is Linked to CA3 Hippocampal GABAergic Integrity

**DOI:** 10.1101/2025.10.18.682786

**Authors:** Cristóbal Ibaceta-González, David Neira, Nicolás M. Ardiles, Nicol Baeza-Araya, Alfredo Kirkwood, Pablo Moya, Adrián G Palacios

## Abstract

The preservation of cognitive function during aging remains a key challenge in neuroscience. In this study, we applied an integrative approach, combining behavioral assays with neurophysiological recordings, to investigate hippocampal circuit integrity. We used *Octodon degus*, a rodent with exceptional longevity (up to 10 years in laboratory conditions), as a natural model of aging and neurodegenerative disease such as Alzheimer. To assess agerelated cognitive changes, we employed three behavioral tasks: Novel Object Recognition (NOR), Open Field (OF), and the Burrowing Test (BT). The BT reflects Activities of Daily Living (ADLs) and is based on species-typical spontaneous burrowing behavior, which has been linked to neurodegenerative markers in degus. We also performed multielectrode electrophysiological recordings to assess GABAergic function in the hippocampus. Aged degus with high BT performance (classified as good burrowers, or GB) showed robust hippocampal activity, especially in the CA3 region, a key hub for signal integration and memory encoding. In contrast, degus with poor BT performance (bad burrowers, or BB) exhibited reduced spontaneous hippocampal activity, suggesting potential compensation via GABA-independent synaptic mechanisms. Altogether, our findings suggest that preserved GABAergic function supports cognitive resilience in aging degus. These results offer new insights into the neural mechanisms underlying healthy cognitive aging and may inform future strategies for preventing or mitigating neurodegeneration.

## Introduction

Aging in the brain introduces numerous biochemical changes that affect neuronal function, leading to reduced synaptic transmission efficiency, altered circuit dynamics, and impairments in signal processing, learning, memory, and behavior (Sibille, 2013). In Alzheimer’s disease (AD) models, hyperactive neuronal states have been observed, particularly in animals exhibiting cognitive deficits (Haberman et al., 2017; Palop et al., 2007). This increased excitability, often due to elevated spontaneous firing in pyramidal neurons, is linked to reduced inhibitory post-synaptic potentials and decreased amplitude and frequency of inhibitory currents. These disruptions alter the excitation/inhibition (E/I) balance and are closely tied to GABAergic signaling via GABA receptors and interneurons (McQuail et al., 2015). Moreover, aging is often accompanied by a reduction in GABA receptor density (by 5–35%), compromising fast inhibitory signaling and weakening inhibitory control (Shen et al., 2010). However, some aged rats that preserve cognitive abilities display enhanced inhibitory synaptic strength in CA1 via tonic inhibition, while maintaining fast inhibition in the dentate gyrus (DG) (Tran et al., 2018). These compensatory mechanisms differ from those in younger animals, which typically do not exhibit such inhibitory adaptations. Supporting this, cognitively intact aged rats show E/I ratios in the DG similar to those of young rats, suggesting preservation of feedforward inhibition through interneuron recruitment (Tran et al., 2019). Together, these findings highlight the importance of maintaining E/I balance, particularly through GABAergic mechanisms, for sustaining cognitive function in aging.

We selected *Octodon degus*, a diurnal, social rodent endemic to Chile, as our model organism, given its natural longevity (8–10 years in laboratory settings) and susceptibility to age-related cognitive decline (Hurley et al., 2018) (Ardiles et al., 2013). Prior studies have shown that aged degus accumulate Alzheimer’s-like pathologies, including amyloid-β (Aβ) and tau aggregates, and exhibit impairments in cognitive tasks such as the T-maze and Novel Object Recognition (NOR) (Inestrosa et al., 2005) (Ardiles et al., 2012). These deficits correlate with synaptic plasticity impairments, including reduced long-term potentiation (LTP) and depression (LTD). However, traditional cognitive assays like the NOR and T-maze do not reliably distinguish between different cognitive trajectories in aged animals, potentially due to task complexity or motivational demands.

To better assess cognitive function in aging, we incorporated the Burrowing Task (BT), a behavior rooted in naturalistic, goal-directed digging activity that parallels human Activities of Daily Living (ADL) (Deacon, 2009)(Dudek et al., 1983). ADLs are among the first functions to decline in neurodegenerative diseases such as AD (Reisberg et al., 2001). Laboratory models have shown that hippocampal damage reduces BT performance (Deacon et al., 2002), and performance in aged degus correlates with AD biomarkers including APP, ApoE, β-amyloid, and neuroinflammatory markers (Deacon et al., 2015). In this study, we aimed to investigate the variability in cognitive aging among degus. We evaluated behavior using the NOR, OF, and BT paradigms, and assessed hippocampal excitation/inhibition (E/I) balance through multielectrode recordings and pharmacological manipulation of the GABAergic system.

## Material and Methods

### Animals

Degus were housed at the animal facility of the Universidad de Valparaíso under controlled conditions (25 °C, 12:12 h light/dark cycle) with ad libitum access to food and water. To minimize stress from handling and transportation, animals were habituated for 45 minutes near the behavioral testing room prior to each session. All behavioral procedures were conducted in the morning (9:00–11:00 a.m.) to control for circadian influences. All experimental procedures were approved by the Universidad de Valparaíso Bioethics Committee (#BEA 141-19) and adhered to international and ANID ethical and biosafety guidelines.

For electrophysiological recordings, degus were anesthetized with isoflurane (3.5% in 600 mL/min O₂; RWD, China) using a dedicated anesthesia chamber (RWD 510) and then euthanized by decapitation. Brains were rapidly extracted, and 300 μm-thick hippocampal slices were obtained using a vibratome (1000 Plus, Vibratome) in ice-cold dissection buffer (4 °C). The dissection buffer contained (in mM): 2.6 KCl, 1.23 NaH₂PO₄, 26 NaHCO₃, 212.7 sucrose, 10 glucose, 3 MgCl₂, and 1 CaCl₂, bubbled with 95% O₂ and 5% CO₂. Slices were then incubated for 1 hour at 32 °C in artificial cerebrospinal fluid (aCSF) composed of 2.6 KCl, 1.23 NaH₂PO₄, 26 NaHCO₃, 124 NaCl, 10 glucose, 1 mM MgCl₂ (adjusted to 3 mM during MEA recordings), and 2 mM CaCl₂, also bubbled with 95% O₂ and 5% CO₂.

### Behavior tests

#### Open field (OF)

Locomotor activity and general exploration were assessed using a circular open field (OF) arena (diameter: 180 cm; wall height: 80 cm) made of white Plexiglas. Each animal was allowed to explore the arena freely for 5 minutes. The surface was cleaned with 70% ethanol between trials to remove odor cues. Behavioral tracking was performed via video recording and analyzed using ANY-maze software. Time spent in the center versus periphery was calculated as a ratio: The exploration of the center and periphery of the OF was calculated as the ratio: (center time) or (periphery time) / (center time + periphery time).

#### Novel object recognition memory (NOR)

Recognition memory was assessed using a three-phase NOR task: (i) Familiarization: Animals explored two identical objects for 180 seconds. (ii) Retention: After a 2-hour delay in their home cage, the objects were replaced with a novel and a familiar one. (iii) Recognition: Animals explored the object pair again for 180 seconds. A preference index (PI) was calculated based on object selective exploration as PI = NO / NO + FO in seconds. Objects were selected of similar sizes in metal, glass, or plastic with similar size.

#### Burrowing test

The Burrowing Test (BT) was adapted from (Deacon, 2006). A plastic burrow tube (30 cm long, 10.5 cm diameter) was filled with 1300 g of rabbit food pellets (not used as food by degus). Animals were placed in the burrowing apparatus for 1 hour, and the weight of displaced pellets was recorded as an index of burrowing performance. A threshold of 130 g (10% of the total) was used to classify animals as Good Burrowers (GB) or Bad Burrowers (BB), following prior studies (Deacon et al., 2015; Deacon et al., 2002).

#### Electrophysiological recordings

Hippocampal slices were placed onto a 252-electrode multi-electrode array (MEA; Multichannel Systems, Germany) mounted in a plastic O-ring chamber and secured with a dialysis membrane (MWCO 25,000; Spectra). Electrodes were 30 μm in diameter, spaced 200 μm apart, and covered a total area of 3.2 cm. The slices were perfused continuously with oxygenated aCSF (5 mL/min, 95% O₂/5% CO₂) at 33°C using a TCO2 temperature controller and PPS2 perfusion system (Multichannel Systems). Prior to recording, slices were stabilized on the MEA for 10 minutes. Electrophysiological data were acquired using MC/Rack software (Multichannel Systems) at a 20 kHz sampling rate. High-resolution images of each slice were taken for reference using an inverted microscope (Nikon ECLIPSE TE200, 5× objective) with a PCO Pixelfly camera (Germany). Each experiment consisted of two recording phases: 10 minutes of spontaneous activity (baseline), followed by 20 minutes of pharmacological manipulation using picrotoxin (PTX, 100μM), a GABA_A_ receptor antagonist. PTX was prepared by diluting 50μL of 100mM stock solution into 45 mL of aCSF, continuously bubbled with 95% O₂/5% CO₂.

#### Spike Sorting and analysis

After completion of an electrophysiological experiment, a spike sorting procedure was performed to separate neurons as described in (Yger et al., 2018). Briefly, spike sorting consists of five main steps: Filtering: The raw extracellular signals are high pass filtered with a Butterworth filter of order three (cutoff of 300Hz). Whitening: Spurious spatial correlations between nearby recordings electrodes are removed in this step. Clustering is a spike sorting core, where action potentials are given to a single unit (template) and separated from other units. The clustering algorithm is based on classifying spikes according to their similarity and separating them. Fitting: Once the template dictionary is done, the fitting process matches the template in recording and thus reconstructs the signal along the whole recording. Merging: Finally, duplicate templates merge to get valid units or neurons.

The spiking-circus software was run using default parameters, only changing the waveform time detection (from 5ms to 3ms). The configuration file is divided into five main steps: the frequency and filter to use; number of clusters and template per electrode; threshold to detect spikes; or set-up experiments condition (number of electrodes and geometric distribution in MEA matrix, the distance among electrodes, etc.). Once cell sorting was done, a manual supervisor (GUI) was required, using a Matlab version GUI. We selected units that accomplished the following criterion: More than 1000 spikes during the complete recording and less than 1% of spikes-pair violate the refractory period (3ms).

#### Firing, maximum firing rate, Inter-spike Interval (ISI), and Burst Rate

We analyzed neuronal activity across experimental groups using the following metrics: (i) Firing Rate (FR): Calculated as the total number of spikes divided by recording duration (10 min during spontaneous activity [SA]; 20 min during PTX application). (ii) Inter-Spike Interval (ISI): Measured as the time between consecutive spikes. (iii) Burst Rate: Defined using ISI-based criteria, where a burst was identified if three or more spikes occurred within a short, neuron-specific time window, as described by (Chen et al., 2009). Burst length was calculated as the average duration of burst events. We also computed the burst spike proportion, defined as the percentage of total spikes that occurred within burst events. All analyses were performed in MATLAB.

#### Hippocampal zones

Neurons were spatially mapped to hippocampal subregions (CA1, CA3, and dentate gyrus [DG]) based on the physical location of the slice over the MEA grid. On each experimental day, a reference image of the hippocampal slice was captured and aligned with the MEA layout. Using spike sorting metadata, including the electrode of origin, neurons were assigned to specific hippocampal zones to facilitate region-specific analysis.

#### Western blot

Degus were euthanized by decapitation, and brains were rapidly extracted. Hippocampi were dissected and homogenized in ice-cold Tris-HCl lysis buffer (1% Triton X-100, 10 mM Tris-HCl, 300 mM NaCl, 5 mM EDTA, pH 7.4) containing a protease and phosphatase inhibitor cocktail (1%; Thermo Fisher Scientific, USA). Homogenates were incubated on ice for 20 minutes, sonicated, and centrifuged at 14,000 × g for 30 minutes at 4 °C. Supernatants were collected and protein concentrations determined using the Qubit™ Protein Assay Kit (Thermo Fisher Scientific). For SDS-PAGE, 60 μg of protein per sample was loaded onto an 8% polyacrylamide gel. Following electrophoresis, proteins were transferred to either nitrocellulose or PVDF membranes (Amersham GE Healthcare, UK), depending on the target protein. Transfers were performed at 30 V for 16 hours at 4°C. Membranes were blocked with 5% non-fat dry milk in either PBS or TBS (pH 7.4) with 0.05% Tween-20 for 1 hour at room temperature. Primary antibody incubations were carried out overnight at 4°C using the following antibodies (1:1000 unless noted): GluR1, GluR2, PSD-95, GluN2A, GluN2B, NR1 (various vendors), and β-actin (1:20,000; Abcam). After washing, HRP-conjugated secondary antibodies were applied (anti-mouse IgG or anti-rabbit IgG; Abcam). Immunoreactive bands were detected using ECL Select substrate (GE Healthcare), and band intensity was quantified using ImageJ (NIH).

#### Network connectivity

Functional connectivity within hippocampal slices was assessed using pairwise Pearson correlation of spike trains across neuron pairs. Because spike train data are discrete and sparse, raw correlation values tend to be low. To address this, we implemented a significance-based thresholding approach. For each neuron pair, spike trains were randomly shuffled 1,000 times to generate a null distribution of correlation values (i.e., chance-level correlations without temporal structure). Observed correlations exceeding this shuffled threshold were considered significant and assigned a binary value of 1 (i.e., a meaningful connection); all others were set to 0. The final network connectivity metric was computed as the percentage of significant correlations out of all possible neuron pairings within each slice. This method allowed us to compare the relative integrity and density of functional networks across experimental conditions (e.g., spontaneous vs. PTX-evoked activity, GB vs. BB groups).

#### Statistical analysis

A Shapiro-Wilk test was performed to evaluate parametric and non-parametric data distribution to determine the statistical significance difference. Parametric data were applied to a test-T to compare the two groups. A Kolmogorov-Smirnov test was applied to compare two groups and the Klustal-Wallis test to compare multi-groups, both tests to nonparametric data. A Fisher’s exact test was used for contingency data, classified into two types. Statistical analysis was done using Prism software (Graphpad Software Inc) and MATLAB.

## Results

### BT is more sensitive to separating two aged degus populations than NOR

To assess memory performance in degus, we used the NOR test, a well-established behavioral assay that does not require external motivation or reinforcement (Ardiles et al., 2012; Ennaceur & Delacour, 1988; Rivera et al., 2021). The test leverages rodents’ innate tendency to explore novel stimuli over familiar ones (Antunes & Biala, 2012).

We tested 17 degus aged 39 to 82 months using the NOR task (Figure 1A). Based on prior studies (Ardiles et al., 2012), a preference index (PI) of 70% was used as the performance threshold to determine task success. Only 4 out of the 17 animals exceeded this threshold (Figure 1A), while the majority showed no significant difference in exploration time between the novel and familiar objects (Figure 1B). These results align with previous findings showing that degus over 36 months often fail to perform the NOR successfully. Additionally, a slight age-dependent decline in PI was observed (Figure 1C). Notably, NOR performance did not correlate with any Open Field (OF) exploration parameters (Figure 1D), suggesting NOR is not confounded by general locomotor activity or anxiety.

**Figure 1.**
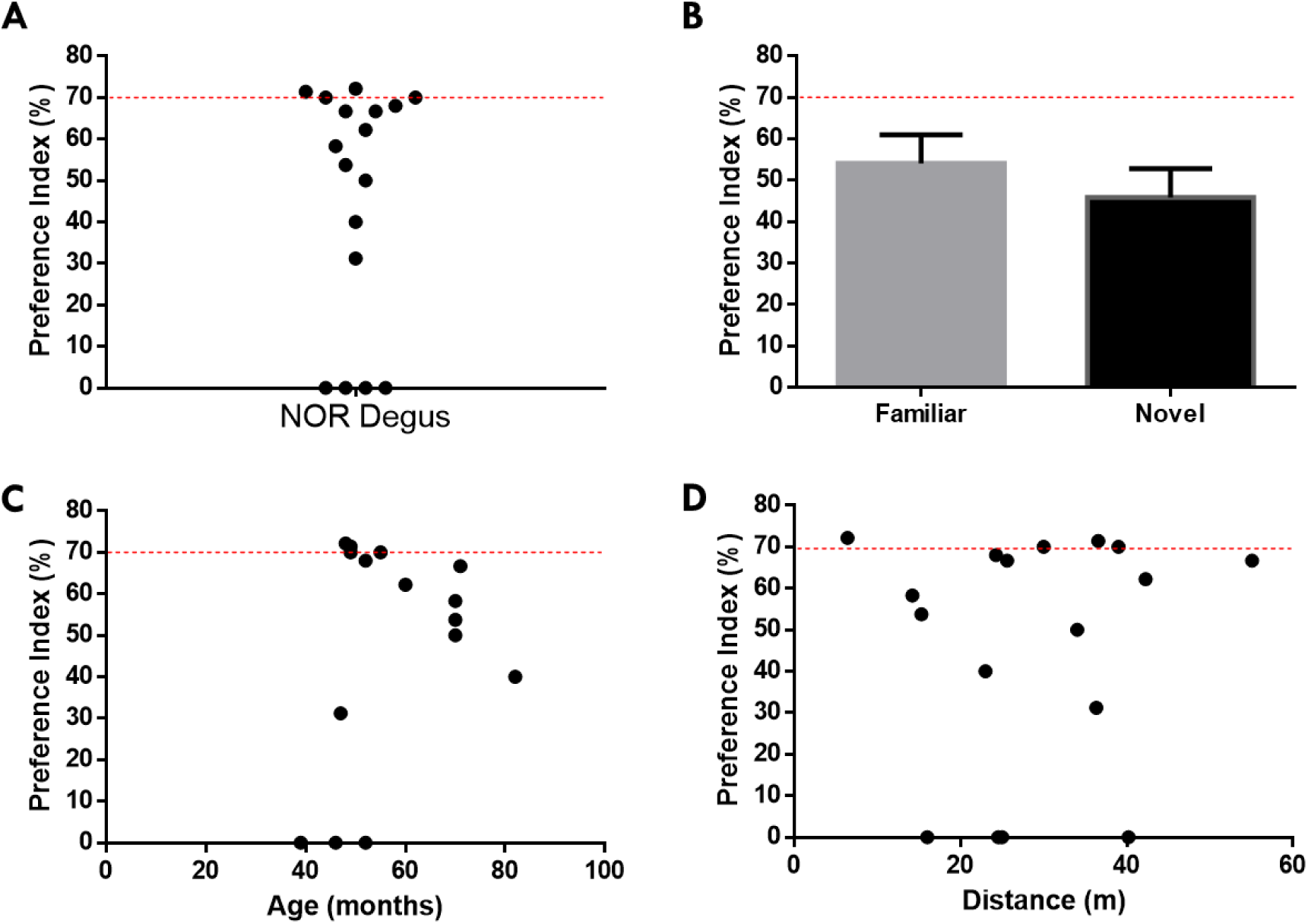
NOR test performance in aged degus. **A** NOR test performance as preference index percentage (n=17). The red line represents the threshold value criterion (70% of the preference index). **B** Comparison between familiar (gray bar) and novel object (black bar) preference index. **C** NOR performance (percentage of preference index) according to degus age (in months). **D** NOR performance (rate of preference index) according to OF performance (distance traveled in meters). Data is shown as mean ± SEM. Statistical analysis using T-test.

We next evaluated burrowing behavior in a separate cohort of 35 degus, aged 24 to 94 months, using the Burrowing Test (BT). Burrowing performance was quantified by weighing the amount of food pellets displaced from the burrow tube (Figure 2A–B). As described in the Methods section, a 10% displacement threshold (130 g of 1300 g total) was used to distinguish Good Burrowers (GB) from Bad Burrowers (BB) (Figure 2B, right).

**Figure 2.**
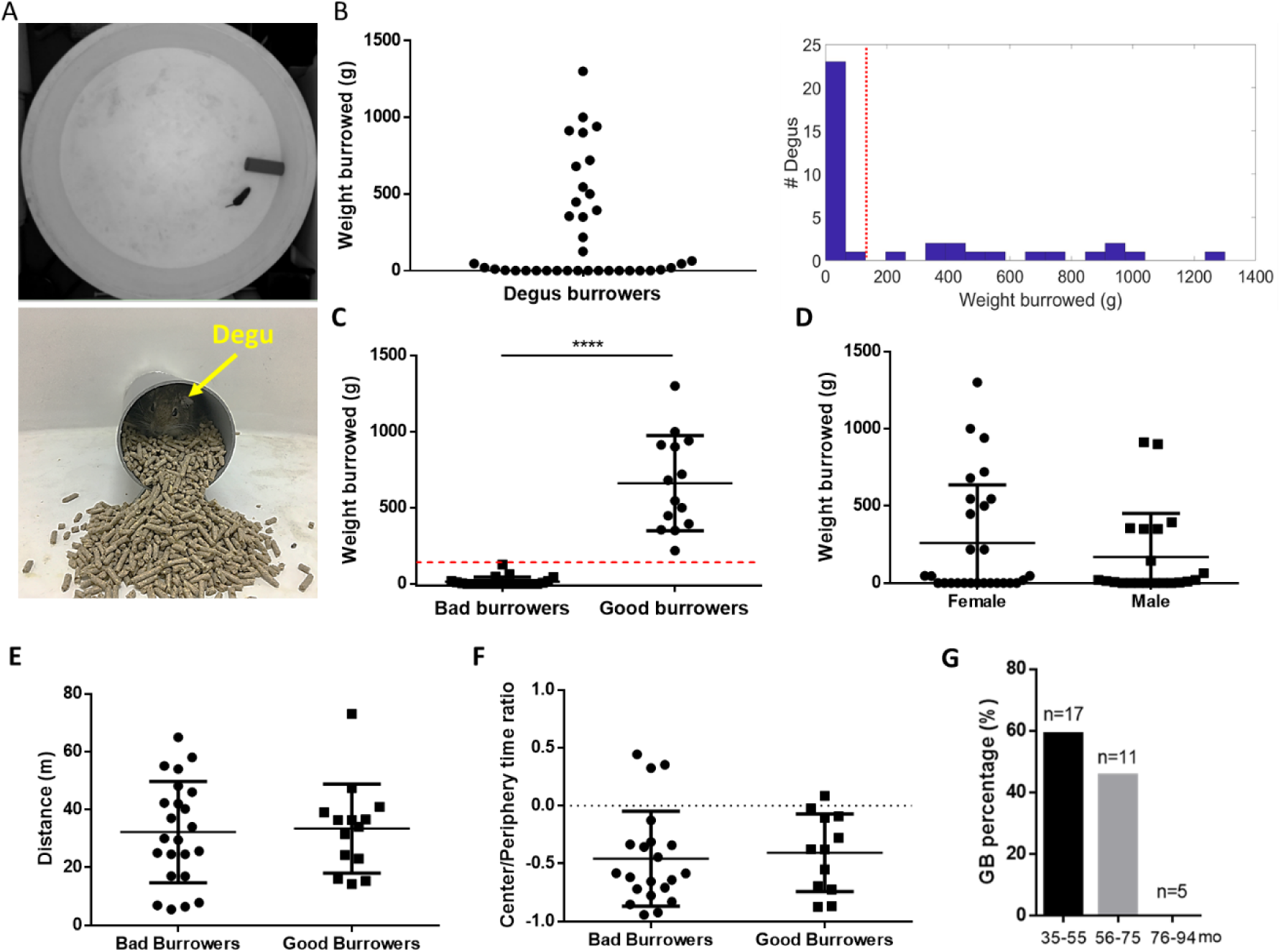
BT measure in aged degus. **A** Localization of the set-up against the wall of a circular OF (diameter 180 cm). The degus were put in the OF for free exploration and BT (top). Degus burrows rabbit food and displace out the content of the burrowing apparatus (bottom). **B** Burrowing performance of degus in terms of weight of pellet burrowed (n=35) (left) and a histogram for selecting a threshold (red line) to split aged degus population (left). **C** Degus were classified according to their performance into good (GB) or bad burrowers BB). According to the histogram in B left, the threshold (red line) value of 10% of the total pellet burrow was determined (130g) to separate GB from BB. **D** Burrowing performance according to degus sex. **E** Traveled distance by degus in OF maze (the same maze of BT) and separated by Bad or Good burrowers. **F** Center-periphery time ratio where degus spend the time (5 min) in the maze, dividing this into two areas, proportionally equal. **G** GB performance divided our colony into three groups with the same time interval (20 months): 35-55 months (black bar), 56-75 months (gray bar), and 76-94 months (no bar). Data is shown as mean ± SD. Statistical analysis using T-test. ****=p<0.0001.

The results showed that 13 degus of 35, or 37.1%, exceeded the 10% threshold and were classified as GB. The remaining 22 (62.9%) were classified as BB and did not exceed the 10% value. Both groups are statistically different (Fig 2C; BB = 15.7 + 33.6 g, GB = 661.9 + 312.4 g; p<0.0001), showing the sensitivity of BT to separate two populations of aged degus, contrary to NOR. Our results did not show bias by sex, where 21 degus were female and 14 males, being statistically similar both groups, according to their BT performance (Fig 2D; females (n=21) = 293.9 + 405.8 g, males (n=14) = 211.6 + 316.6 g; p = 0.49).

To rule out confounding effects of mobility or anxiety during BT, we analyzed OF behavior during the first 5 minutes of burrowing trials. There was no significant difference in total distance travel between GB and BB groups (GB = 33.4 ± 15.4 m; BB = 32.2 ± 17.5 m; *p* = 0.84; Figure 2E). Similarly, the center-to-periphery exploration ratio was comparable (GB = – 0.41 ± 0.34; BB = –0.46 ± 0.41; *p* = 0.72; Figure 2F), indicating both groups preferred the periphery—consistent with established exploratory behavior in degus (Ardiles et al., 2012; Rivera et al., 2021).

We further stratified the degus into three age groups: 35–55, 56–75, and 76–94 months (Figure 2G). A clear age-dependent decline in BT performance was observed. In the 56–75– month group, 45.5% of animals were classified as GB, while in the oldest group (76–94 months), no animals met the GB criteria. This trend reinforces the idea that BT performance, analogous to Activities of Daily Living (ADLs), declines with age, similar to early functional losses seen in neurodegenerative conditions such as Alzheimer’s disease AD (Deacon, 2009; Reisberg et al., 2001).

### Hippocampal neurons of GB showed a higher response to PTX than BB

Once behavioral tasks were done, we evaluated the hippocampal network state of BB and GB through electrophysiology technique, specifically interested in GABAergic transmission. Each experiment consists of 10 min of spontaneous activity (SA) recording, followed by 20 min of recording activity induced by the picrotoxin drug (PTX), an allosteric inhibitor of ionotropic GABA receptors (GABA_A_). Figure 3A (top) shows a representative raster plot, each line corresponds to a single neuron, sorted from low to high firing rate for BB (n = 162, left) and GB (n = 143, right). The global activity (Fig 3A bottom) was normalized according to the total number of neurons recorded per group by each bin (1 s). PTX increases neuronal activity in both BB and GB conditions, but GB neurons seem more active than BB. To quantify the effect of the PTX, we select the last 10 min of each recording, representing a steady-state level value of the system (Burke & Barnes, 2006).

**Figure 3.**
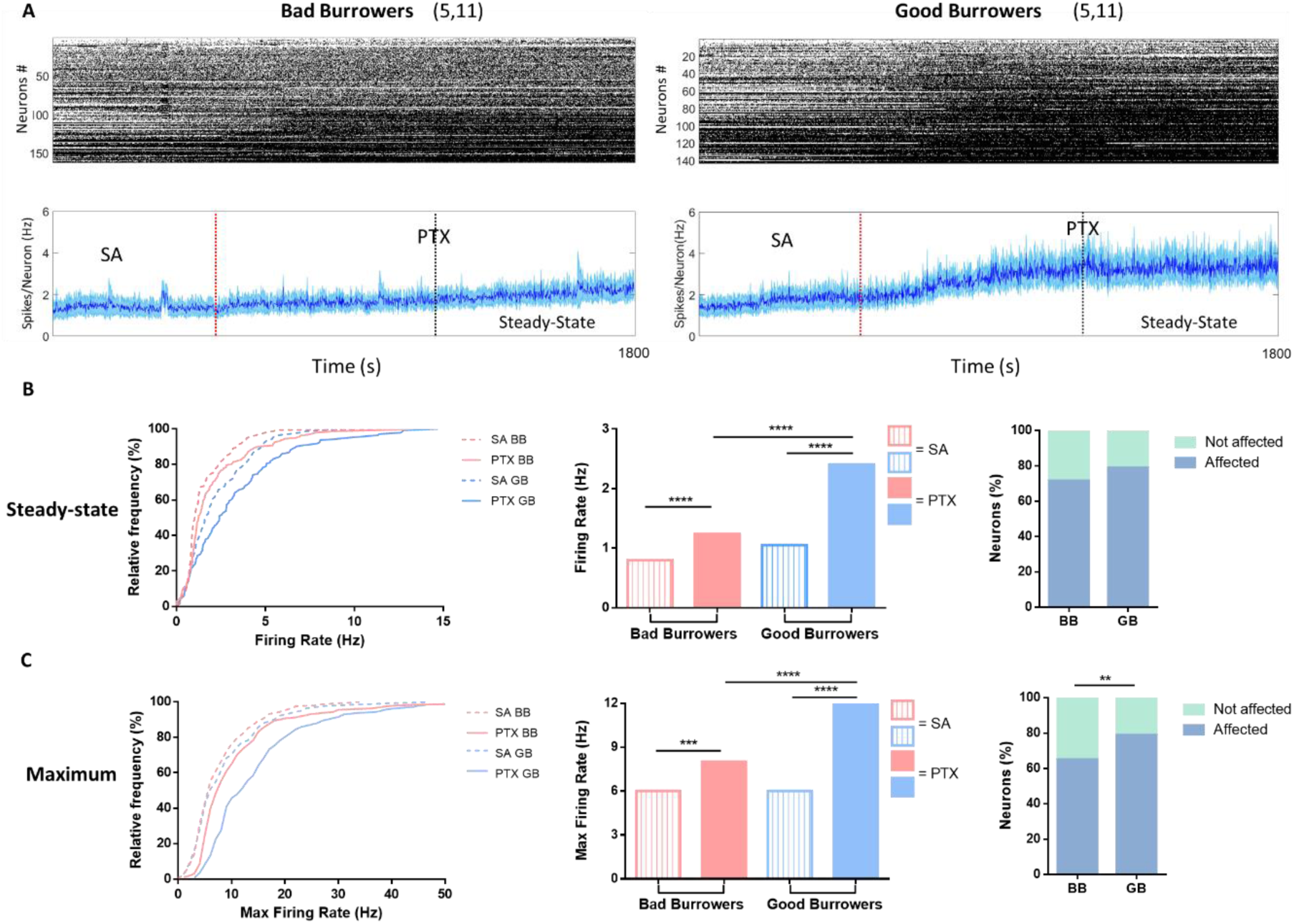
MEA recording for GB and BB degus. **A** Representative raster plot for whole neurons was recorded, separated according to BT performance into bad (5 degus, 11 slices, n=162 neurons, left) and good (5 degus, 11 slices, n=143, right) burrowers (top). Population activity represented by spikes/neuron (Hz) (blue line) + 95% of confidence error (blue shaded) for the whole recorded time (1800 s). The redline separates spontaneous activity (SA, 600 s, left) and PTX activity (PTX, 1200 s, right). The black line represents the last 600 s where was compute the steady-state firing rate (bottom). Bin selected was 1s. **B** SA (segmented line) and PTX (continue line) steady-state firing rate (FR) separated into BB (pink) or GB (blue), represented by cumulative frequency plot (left). The median of FR cumulative frequency distribution for SA (empty bar with lines) and PTX (filled bar), separated into BB (pink) or GB (blue) (center). The number of neurons affected by PTX (calypso blue) increasing their SA FR > 1 after PTX induction (steady state FR) (right). In Hertz, **C** Maximum FR activity is like B, using a bin size of 1s. Data showed a median. Statistical nonparametric test Kolmogorov-Smirnov and Fisher’s exact test for affected and not affected values. **=p<0.01; ***=p<0.001; ****=p<0.0001. Both rasters were sorted increasingly, according to whole neuronal FR.

A more quantitative measure was carried out comparing the firing rate (FR) (Fig 3B, left) and Maximum FR (Fig 3C, left). The BB and GB increased neuronal activity under PTX, both during the steady state FR and during the maximum FR, compared with their SA (FR BB: SA = 0.81 Hz; PTX = 1.2 Hz; p<0.0001. GB: SA = 1.16 Hz; PTX = 2.41 Hz; p<0.0001) (Maximum FR BB: SA = 6 Hz; PTX = 8 Hz; p<0.001. GB: SA = 6 Hz; PTX = 12 Hz; p<0.0001) (Table 1). This was expected because a drug that inhibits the GABAergic system should raise neuronal activity.

Since the data were non-parametric, values are represented as medians and analyzed using cumulative frequency plots and appropriate statistical tests (e.g., Kolmogorov–Smirnov).

When we compare the Maximum FR for GB and BB, they show an acute effect of PTX. This is consistent with the response that PTX elicits over the hippocampal slice, generating epileptic-like responses as seizures due to a disruption of E/I balance (Hablitz, 1984; Hashimoto et al., 2017).

An important result here was the observation that BB degus showed a tendency for less SA activity compared to GB degus, and we interpret it as the presence of a compensatory mechanism different from the GABAergic system controlling glutamatergic activity.

To quantify the number of neurons affected by PTX, we separated their responses based on their activity during SA. The separation consisted of identifying those that increased (affected) their activity with PTX from those that decreased (not affected) it (Fig 3B-C, right). The FR during steady state and maximum FR showed a higher number of neurons involved by PTX (FR: BB = 69.75%; GB = 72.72%; p>0.05. Max FR BB = 65.0%; GB = 78.9%; p<0.01.), where the difference on maximum FR was significant according to Fisher’s exact test, supporting the idea that GB tends to preserve more neurons modulated by the GABAergic system.

Burst is “a group of action potentials generated in rapid succession, followed by a period of relative quiescence” (Zeldenrust et al., 2018). The functional meaning of burst is related to plasticity modulation, enhanced reliability in information transmission, and expanding the coding space. We used an algorithm based on ISI distribution, defining a burst detector threshold for each neuron (Chen et al., 2009), and it has been used in different works (Bridi et al., 2018; Shin et al., 2021; Trujillo et al., 2019). This threshold is a time window whereby at least three spikes consider a burst event.

Figure 4A shows the burst activity (left and center) and the number of affected neurons by PTX (right). Like FR, the burst activity increased under PTX activity on both BB (SA = 0.016 Hz; PTX = 0.020 Hz; p<0.01) and GB (SA = 0.047 Hz; PTX = 0.086 Hz; p<0.0001). Comparing the burst distribution under PTX activity, we find that the burst activity was higher for GB than BB (p<0.0001). The number of neurons affected by PTX also showed a significant sensibility for PTX (BB = 63.8%; GB: = 77.5%; p<0.01).

**Figure 4.**
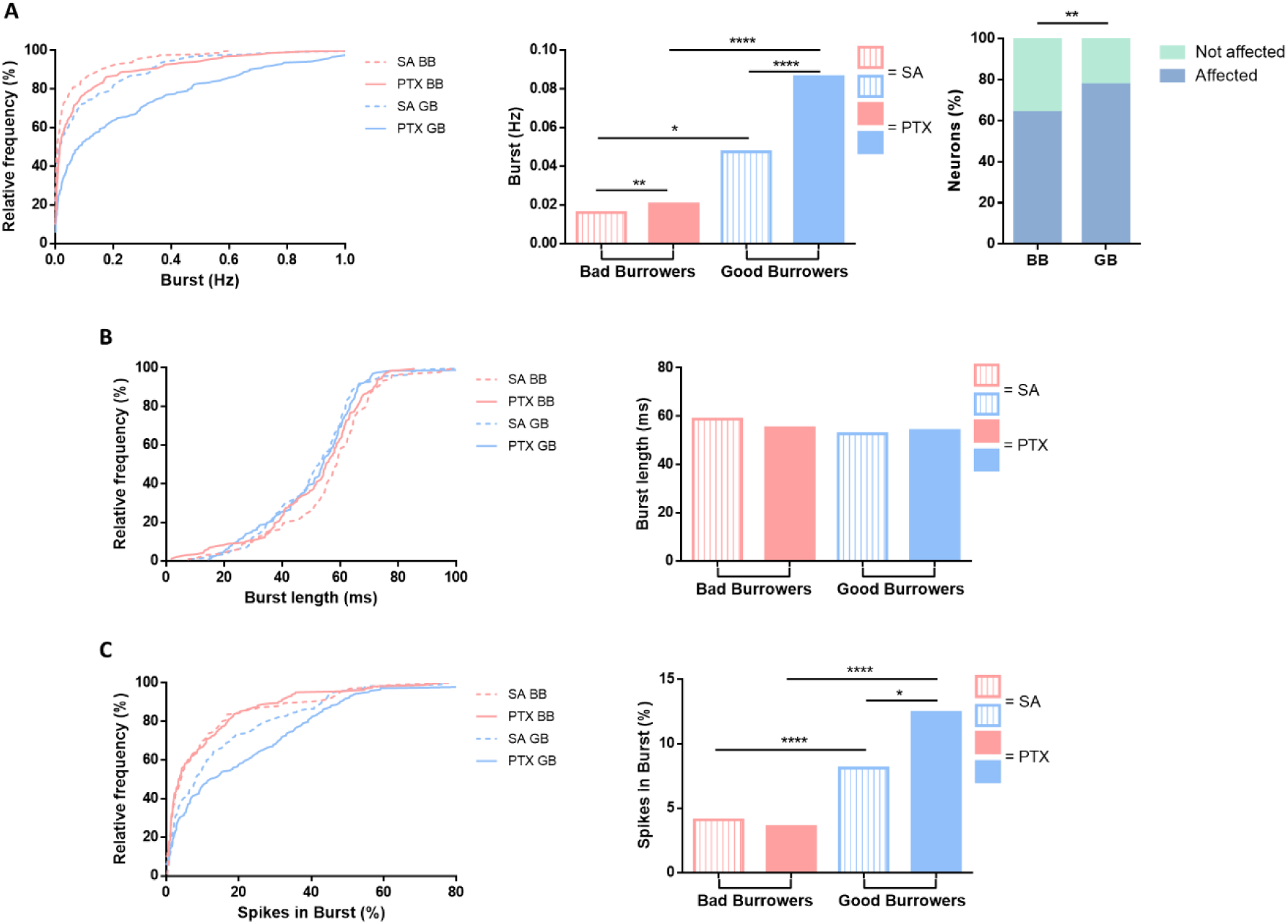
PTX and burst activity. **A** Burst activity, in Hertz, for SA (segmented line) and PTX (continue line) steady state, separated into bad (BB; pink) or good burrowers (GB; blue), represented by cumulative frequency plot (left). The median of burst cumulative frequency distribution for SA (empty bar with lines) and PTX (filled bar), separated into BB (pink) or GB (blue) (center). The percentage of neurons affected by PTX (calypso blue), increasing more than one time their SA burst on PTX activity (steady state FR) (right). **B** Cumulative frequency plot of burst length (ms, left) and percentage of spikes that belong at a burst (**C**), using the same representation on A. Data shows the median. Statistical nonparametric test Kolmogorov-Smirnov and Fisher’s exact test for affected and not affected values. *=p<0.05; **=p<0.01; ****=p<0.0001.

This data strongly suggests that the GABAergic system of GB tends to be more responsible and in better health condition than BB. According to previous descriptions, the GABA ionotropic circuits are affected during aging, losing their functional control of neuronal circuits and, therefore, their E/I balance (Rozycka & Liguz-Lecznar, 2017). The correct E/I balance has been described as a critical mechanism for preserving cognitive skills during aging (Tran et al., 2019; Tran et al., 2018). Wherein GB degus performing better in BT suggest that their GABAergic systems are better maintained than BB. Moreover, BB showed a lower SA during steady-state FR and Burst needs some explanation. If BBs were to control only their excitatory activity via the GABAergic system, we would expect to see a high SA in line with work that has linked aging with hyperactivity (Haberman et al., 2017).

Figure 4 shows the burst length (B) and spikes contained in bursts (C). This experiment did not find a difference among all groups (BB: SA = 58.8 ms; PTX = 55.2 ms; p=0.202. GB: SA = 52.7 ms; PTX = 54.1 ms; p=0.256). Furthermore, spikes in burst increase under PTX effect but only for GB (BB: SA = 4.09%; PTX = 3.6%; p=0.964. GB: SA = 8.13; PTX = 12.4; p<0.05).

Our results suggest a change in the distribution of spikes in GB, where PTX, besides increasing spikes, also affects the pattern structure of the burst. BB, despite increasing its activity with PTX, its spikes do not follow changes in the burst pattern structure, which is an argument to support the idea that BB does not have a healthy hippocampus (Hablitz, 1984). In conclusion, GB responds to PTX more strongly than BB, acutely (maximum FR), and according to the PTX effect (bursting activity).

### CA3 hippocampal regions of GB are the most affected area by PTX

We used a hippocampal reconstruction map to separate our neurons according to hippocampal zones to contrast the recording electrodes and their spike response in each hippocampal slice. Figure 5A shows a picture of the hippocampal slice over our MEA matrix (left) and a reconstruction map (right), where blue circles represent neurons. Some blue circles are outside the picture because the camera field does not cover the MEA matrix surface.

**Figure 5.**
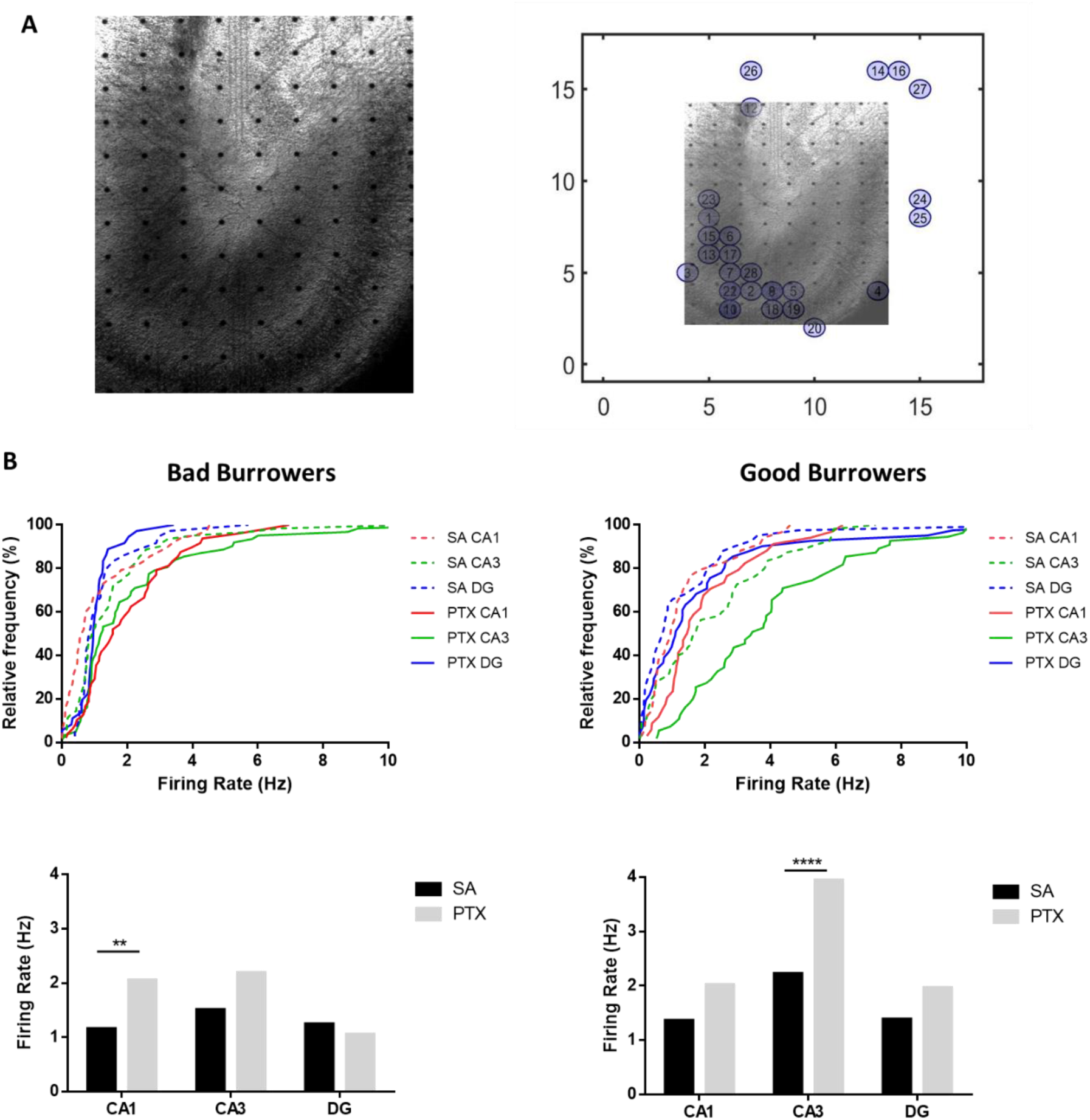
Hippocampal classification according to CA1, CA3, and DG zones. **A** Hippocampal slice over MEA matrix picture in our recording set-up (left). Hippocampal map reconstruction using neurons recorded and successfully sorted (blue circle) compared with a picture of hippocampal slice on MEA matrix (left) **B** Firing rates (FR) to each hippocampal zone (CA1 red, CA3 green and DG blue) according to their burrowing performance (BB left or GB right) and different activity (SA segmented line, PTX continue line), represented by cumulative frequency plot (top). The median of FR cumulative frequency distribution for SA (black bar) and PTX (gray), separated into BB (left) or GB (right) (bottom). Data shows the median. Statistical nonparametric test Kolmogorov-Smirnov. **=p<0.01; ****=p<0.0001.

Once the neurons were localized and classified in a hippocampal zone, their FR of BB and GB in SA and PTX activity were computed (Fig 5B). The results show that BB increased FR under the PTX effect only in the CA1 region, compared with SA (p<0.01) (Table 2). In contrast, GB increased FR only in the CA3 region (p<0.0001) but in a higher significant difference than BB CA1.

Besides finding different responses to PTX for BB and GB hippocampus, we also found differences depending on the hippocampal zone. First, DG showed less activity in both groups, and this can be explained since where granular cells have lower firing rates, which may be related to their participation in sparse coding for pattern separation (Diamantaki et al., 2016; Neunuebel & Knierim, 2012), and being DG the most silent zone of the hippocampal network (Jung & McNaughton, 1993). Our findings are concord with the literature and support our electrophysiology MEA recording.

Our results also show an essential inhibition for CA1 in BB degus and CA3 in GB degus. As mentioned, the E/I balance is a critical mechanism to preserve cognition during aging. Aged rats with similar cognitive skills compared to young rats tend to have a similar E/I ratio (small ratio), predominating inhibition (Tran et al., 2019; Tran et al., 2018). However, the E/I balance is a complex and delicate subject with some contradictory results. For example, some works have shown cognition impairment with the highest CA1 inhibition and LTP decreases (Chapman et al., 1998; Cunha et al., 2019) and spatial memory deficit (Valbuena et al., 2019). We have shown that aged degus showed a deficit in LTP and cognition impairment (Ardiles et al., 2012), which can be an orientation for further studies on SP according to degus BT performance.

The CA3 is the core of the neural circuits of the hippocampus. We found the highest difference compared to GB and BB (maximum FR, number of neurons affected by PTX, Burts activity) (Fig 6, Table 3). Figure 6A shows a representative raster plot of CA3 neurons using the parameters in Figure 3A (BB left and GB right). We observe that PTX induces an increase of the maximum FR for GB (SA = 5 Hz; PTX = 10 Hz; p <0.001) but not for BB (SA = 6 Hz; PTX = 6 Hz; p = 0.982) (Fig 6B). A similar observation for the maximum FR for both groups for SA (BB vs. GB), differing from the observation for whole hippocampus steady-state FR.

**Figure 6.**
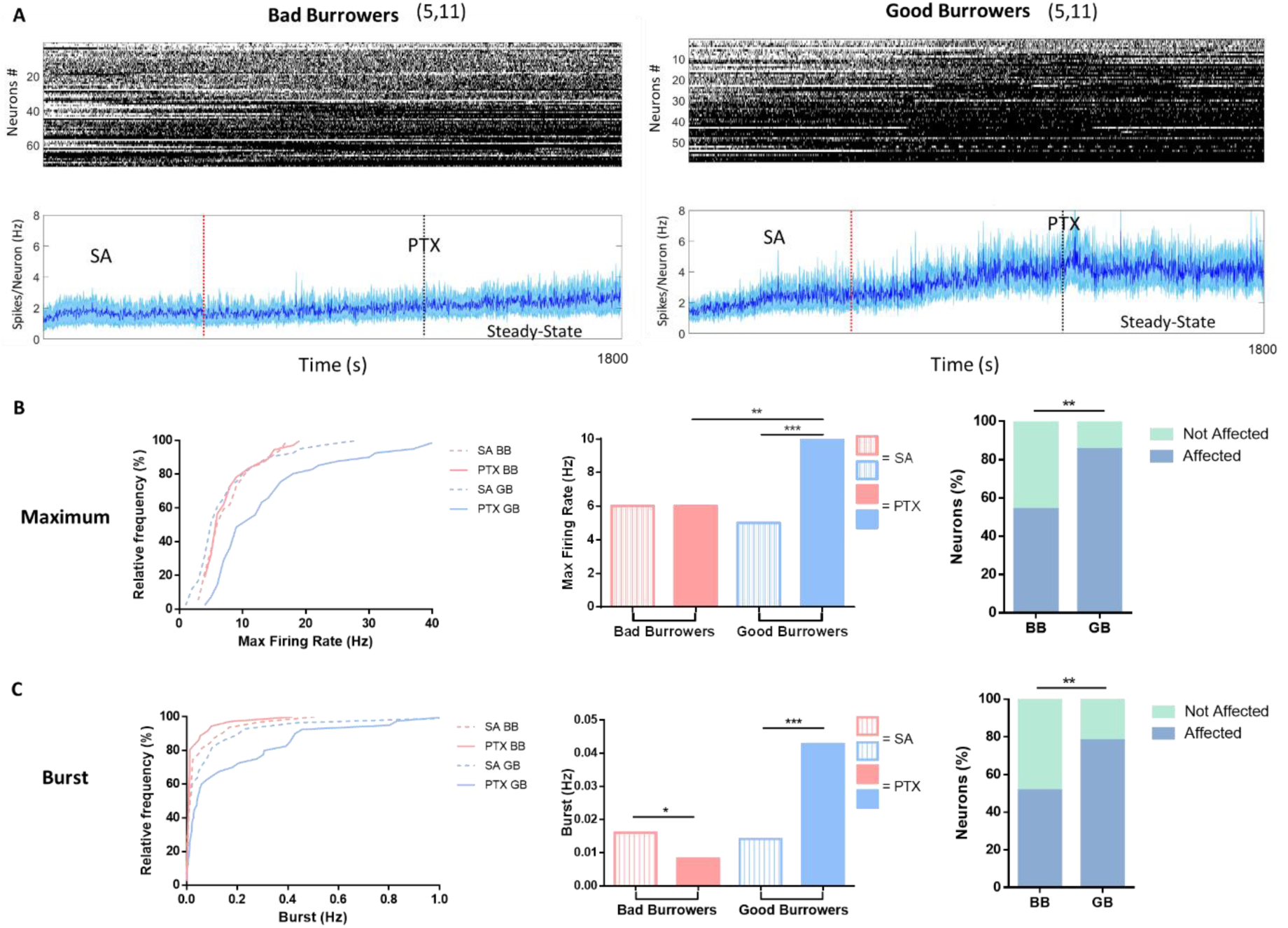
CA3 and PTX. **A** Representative raster plot for CA3 neurons was recorded, separated according to BT performance into bad (n=62 neurons, left) and good (n=55, right) burrowers (top). Population activity represented by spikes/neuron (Hz) (blue line) + 95% of confidence error (blue shaded) for the whole recorded time (1800 s). The red line separates spontaneous activity (SA, 600 s, left) and PTX activity (PTX, 1200 s, right). The black line represents the last 600 s where was compute the steady-state firing rate (bottom). Bin selected was 1s. **B** SA (segmented line) and PTX (continue line) maximum FR on steady state, separated into bad (BB; pink) or good burrowers (GB; blue), represented by cumulative frequency plot (left). The median of maximum FR cumulative frequency distribution for SA (empty bar with lines) and PTX (filled bar), separated into BB (pink) or GB (blue) (center). Neurons affected by PTX (calypso blue), increasing than one time their SA maximum FR on PTX activity (steady state maximum FR) (right). **C** Burst activity, in Hertz, with the same representation in B. Data shows the median. Statistical nonparametric test Kolmogorov-Smirnov and Fisher’s exact test for affected and not affected values. *=p<0.05; **=p<0.01; ***=p<0.001.

The number of neurons affected by PTX was higher for GB than BB (GB = 83.3%, BB = 54.1%; p<0.01). Moreover, the numbers of neurons increasing their maximum FR by > 2 were 40.5% for GB and 8.1% for BB (p<0.001). We also observe that the burst activity for the GB increased under the PTX effect (SA = 0.014 PTX = 0.042; p<0.001) and the neurons affected (BB = 45.2%; GB = 70.4%; p<0.01).

In conclusion, the evidence here shows a significant difference in GABAergic state between BB and GB in the CA3 region, compared with differences observed in the whole hippocampus. Our findings are consistent with previous reports regarding the effect of PTX and the induction of epileptic responses for CA3 (Hablitz, 1984). Indeed, CA3 pyramidal neurons are a pacemaker of spontaneous burst activity, eliciting this specific activity (Witter, 2007). Inhibition is crucial in maintaining quiet CA3 neurons through GABA_A_ receptors (Aradi & Maccaferri, 2004) and, hence, the CA3 network. Our observations support the significant difference between BB and GB under the PTX effect, based on a differential inhibition of the GABAergic system, which is more prominent for CA3, and we speculate on the importance of the CA3 region to maintain a better performance in our aged GB degus preserving part of some of their cognitive skills.

### Network connectivity better preserve in GB

The neural coding population activity is critical to establish learning, memories, and cognition (Doiron et al., 2016). In this context, we assessed a neural population analysis to unveil the hippocampal network state in our GB and BB degus population.

We use the most straightforward network analysis based on pairwise Pearson correlation. However, since the data are discrete, Pearson values were relatively small; hence we introduce a method to find the relevant correlations. It consists of taking the spikes of neurons and shuffling, disrupting the time influences, and computing Pearson correlation, repeating that process 1000 times. All correlation values higher than the “shuffle correlation threshold” (chance threshold) were considered as 1 (relevant connection). Finally, we computed each neuron’s relevant relationships (network connectivity) in the recorded slice. We took the maximum possible correlation of pairwise neurons and divided the sum of relevant connections per neuron.

Figure 7A shows similar network connectivity for BB in SA (16.7%) and PTX (19.1%) (p = 0.054). GB increased network connectivity under the PTX effect (SA = 11.1%; PTX = 28.6%. p<0.0001) more than twice. The GB degus would show a neural network with fewer connections compared to BB in SA (p<0.0001), even if GB showed higher FR (Fig 3B). Nevertheless, when we reached both groups under the PTX effect, GB increased to higher network connectivity than BB (p = 0.0001), reversing previous differences.

**Figure 7.**
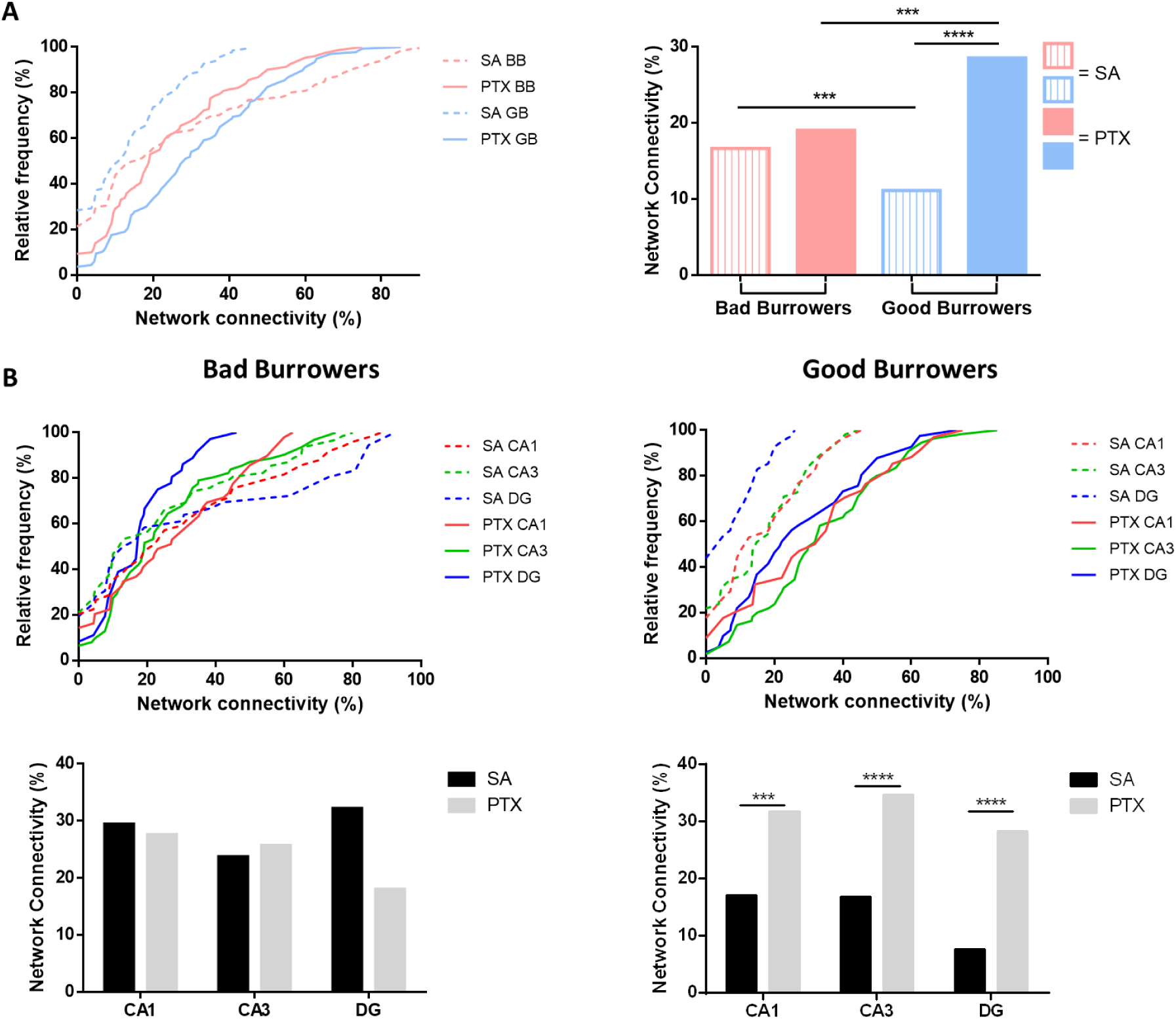
Network connectivity under PTX effect. Network connectivity, computing through Pearson correlation, in spontaneous activity (SA segmented line) and PTX drug (continues line), separated in bad (pink) and good (blue) burrowers, represented by cumulative frequency plot (left). The median of network connectivity cumulative frequency distribution for SA (empty bar with lines) and PTX (filled bar), separated into BB (pink) or GB (blue) (right). **B** Network connectivity to each hippocampal zone (CA1 red, CA3 green, and DG blue) according to their burrowing performance (BB left or GB right) and different activity (SA segmented line, PTX continue line), represented by cumulative frequency plot (top). Median of network connectivity cumulative frequency distribution for SA (black bar) and PTX (gray), separated into BB (left) or GB (right) (bottom). Data are shown as box plot and median + Tukey test distribution. The black circle represents the out-layers. Statistical nonparametric test Kolmogorov-Smirnov. ***=p<0.001; ****=p<0.0001.

Our network connectivity analysis by hippocampal zones showed different results depending on the degus group. BB did not show a significant difference between SA and PTX among hippocampal zones (CA1 SA = 22.2%; PTX = 29.9%; p=0.708. CA3 SA = 13.0%; PTX = 19.2%; p=0.614. DG SA = 14.9%; PTX = 17.4%; p=0.020) (Fig 7B, left). CA1 region has few associational connections, being a network focused on smaller cell modules, although it does not mean that CA1 lacks these connections (Yang et al., 2014). This feature of CA1 could explain why it did not increase its network connectivity, although neural activity was affected by PTX. Consistent with our finding, a network with few recurrent connections (over them self-inputs) is disabled to increase correlation with the network’s neurons.

In terms of connectivity, the GB degus showed significant differences in all hippocampal zones (CA1 SA = 12.5%; PTX = 32.6%; p=0.0013. CA3 SA = 14.8%; PTX = 31.8%; p<0.0001. DG SA = 5.0%; PTX = 22.2%; p<0.0001) (Fig 7B, right). The FR, maximum FR, and burst activity were increased by PTX only in CA3 (Fig 3B and Fig 4B-C), where network connectivity increases in every zone, including DG, where connections from CA3 to DG are almost null. One of the main features that can explain our findings is the recurrent architecture of the CA3 network.

## Discussion

In this study, we combined behavioral, electrophysiological, and computational approaches to investigate the relationship between cognitive health and the integrity of the GABAergic system in aging *Octodon degus*. We introduced the burrowing task (BT), a spontaneous behavior analogous to activities of daily living (ADL), to classify aged degus into good burrowers (GB) and bad burrowers (BB). GB animals showed better preservation of hippocampal GABAergic function than BBs, as evidenced by higher firing rates (Figure 3) and greater burst activity (Figure 4) following pharmacological inhibition of GABA receptors using picrotoxin (PTX). The CA3 region was the most affected in BB animals, showing significantly fewer PTX-responsive neurons in terms of both maximum firing rate and burst activity (Figure 6). Additionally, GB animals exhibited lower network connectivity during spontaneous activity (SA), but higher connectivity under PTX in all hippocampal subregions (Figure 7). Collectively, our results point to a strong correlation between BT performance and GABAergic system integrity, with CA3 emerging as the most preserved region in GB animals (Figure 8).

**Figure 8.**
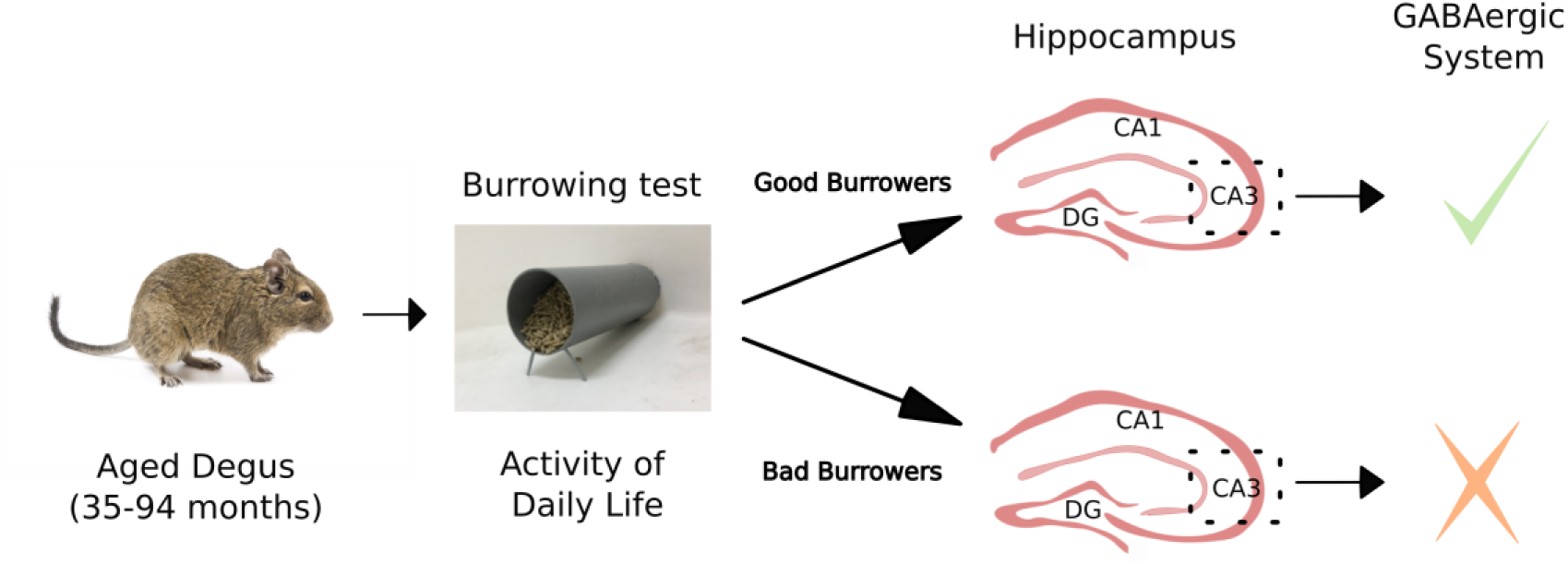
Burrowing test, an Activity of daily life test, allowed us to separate into two populations our aged degus. According to their performance we classified them as good or bad burrowers. Measuring their hippocampal neuronal activity, in presence of a GABA_A_ receptor blocker (PTX), good burrowers had a preserved GABAergic system, specifically on CA3 region.

Selecting an appropriate behavioral test is essential for evaluating the cognitive status of animals. The NOR test is widely accepted as a reliable measure of cognitive performance in *Octodon degus* (Ardiles et al., 2012; Lindsay et al., 2020; Rivera et al., 2021). However, previous studies have shown that degus older than 24 months often display reduced performance on this task (Ardiles et al., 2012), a result that we also observed in our cohort (Figure 1). Importantly, this does not necessarily imply that the animals are cognitively non-functional. Instead, it reflects that NOR performance requires highly preserved cognitive function (Wooden et al., 2021).

In contrast to NOR, the BT allowed us to clearly distinguish between two populations of aged degus over 24 months: GB and BB (Figure 2). To ensure the robustness of this behavioral classification, we included several control measures, such as sex, locomotor activity, and anxiety levels, and found no significant differences between GB and BB animals in these variables (Figure 2). We also assessed the impact of aging on BT performance and observed a progressive decline with age (Figure 2), similar to the age-related deterioration seen with NOR (Figure 1). These findings are consistent with previous reports indicating that ADL-like behaviors, such as burrowing, tend to decline with age (Deacon, 2009; Reisberg et al., 2001).

Our electrophysiological results indicate that GB animals retain a more functional GABAergic system, as demonstrated by their stronger response to PTX application. PTX induces an epileptiform-like effect by transiently increasing neuronal activity, with the CA3 region typically showing the most pronounced response (Hablitz, 1984; Hashimoto et al., 2017). This is consistent with our observations: in GB animals, CA3 neurons were strongly activated by PTX, resembling the behavior of a healthy hippocampal circuit.

The hippocampal CA3 region is not only a computational hub within the hippocampal network (Knierim, 2015), but also one of the most vulnerable areas to seizures and neurodegenerative processes (Cherubini & Miles, 2015). Our results highlight that the key difference between GB and BB animals lies in the functional state of CA3. A well-preserved inhibitory (GABAergic) system in CA3 is critical for maintaining both local circuit stability and the broader hippocampal network that supports cognition and behavior (Aradi & Maccaferri, 2004). The increased network connectivity observed in GB animals under PTX reflects how disinhibition activates the recurrent excitatory circuitry of CA3, synchronizing activity across the hippocampus.

Interestingly, BB animals exhibited lower spontaneous activity (SA) than GB animals, even though they were less responsive to PTX (Figure 3). If BB animals lacked functional GABAergic inhibition, one might expect elevated SA, as reported in other aged animal models (Haberman et al., 2017; Palop et al., 2007). This apparent paradox suggests that BB animals may regulate activity not through inhibition but by suppressing excitation. Glutamate, the primary excitatory neurotransmitter in the central nervous system, is present in over 90% of neurons via various glutamate receptor subtypes (Gasiorowska et al., 2021). Prior research has linked beta-amyloid pathology in AD to deficits in glutamatergic synaptic transmission and retraction of dendritic spines (Palop et al., 2007; Palop et al., 2003). Such alterations may suppress excitatory signaling in a manner distinct from intact GABAergic inhibition.

In line with this, studies in AD transgenic mice have shown that long-term potentiation (LTP) deficits can result from excessive inhibition. In those models, PTX application failed to restore LTP, indicating persistent inhibitory dysfunction (Palop et al., 2007). In our study, we did not detect differences in the expression of glutamatergic receptor subunits (NR1, NR2A, NR2B, GluR1, GluR2, or PSD-95) between BB and GB groups (Supplementary Figure 1). However, because these protein measurements were conducted on whole hippocampal lysates, it is possible that region-specific differences, such as in CA3 or CA1, were masked by averaging across the entire hippocampus.

Overall, our findings support a strong relationship between cognitive performance in aged degus, as measured by the burrowing task, and the integrity of the hippocampal GABAergic system, particularly within the CA3 region. This conclusion is consistent with previous research highlighting CA3 central role in both hippocampal computation and vulnerability to age-related dysfunction.

## Acknowledgments

We thank the Instituto de sistemas complejos for giving us space to do the data analysis. We would also thank Ruben Herzog, Cesar Reyes and Maria José Escobar for their support and assistance with informatics tools.

## Author Contributions

Cristobal Ibaceta-González (Conceptualization; Investigation; Data curation; Formal analysis; Methodology; Writing-original draft; Project administration); David Neira (Investigation); Nicolas M. Ardiles (Investigation; Formal analysis); Nicol Baeza-Araya (Investigation; Formal analysis); Alfredo Kirkwood (Conceptualization; Writing-review and editing); Pablo Moya (Formal Analysis; Writing-review and editing); Adrian Palacios (Conceptualization; Formal analysis; Methodology; Writing-review and editing; Supervision; Project administration)

## Ethical considerations

Protocols for animal experiments were approved by the Universidad de Valparaíso Bioethics Committee (Approval no. #BEA 141-19 on November 08, 2019) and adhered to international and ANID ethical and biosafety guidelines.

## Consent to participate

Not applicable

## Consent for publication

Not applicable

## Supplementary

**Supplementary 1.**
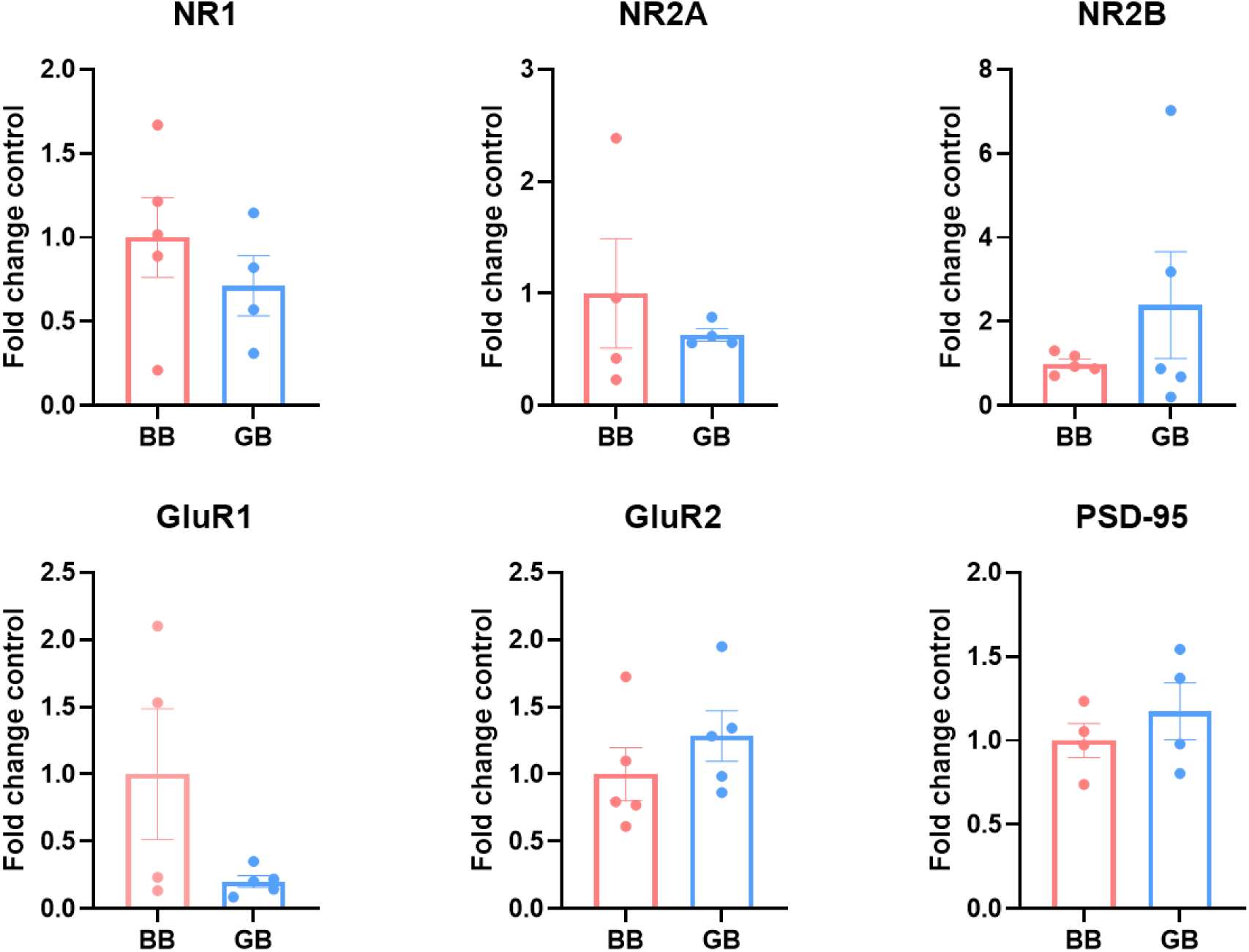
Western blot analysis of different Glutamatergic subunits receptor, such as NR1, NR2A, NR2B, GluR1, GluR2 and PSD-95. Data was normalized using β-actin. Data are shown as mean + SEM. Statistical analysis was T-test.

